# Calcium binding site in AA10 LPMO from *Vibrio cholerae* suggests modulating effects during environment survival and infection

**DOI:** 10.1101/2023.12.22.573012

**Authors:** Mateu Montserrat-Canals, Kaare Bjerregard-Andersen, Henrik Vinter Sørensen, Gabriele Cordara, Gustav Vaaje-Kolstad, Ute Krengel

**Affiliations:** Centre for Molecular Medicine Norway, University of Oslo, NO-0318 Oslo, Norway; Department of Chemistry, University of Oslo, NO-0315 Oslo, Norway; Faculty of Chemistry, Biotechnology and Food Science, Norwegian University of Life Sciences (NMBU), NO-1433 Ås, Norway

## Abstract

Despite major efforts towards its eradication, cholera remains a major health and economic burden in many developing countries. Between outbreaks, the bacterium responsible for the disease, *Vibrio cholerae*, survives in aquatic environmental reservoirs, where it commonly forms biofilms, *e.g.*, on zooplankton. *N*-acetyl glucosamine binding protein A (GbpA) is an adhesin that binds to the chitinaceous surface of zooplankton and breaks its dense crystalline packing thanks to its lytic polysaccharide monooxygenase (LPMO) activity, which provides *V. cholerae* with nutrients. In addition, GbpA is an important colonization factor associated with bacterial pathogenicity, allowing the binding to mucins in the host intestine. Here, we report the discovery of a cation-binding site in proximity of the GbpA active site, which allows Ca^2+^, Mg^2+^ or K^+^ to bind close to its carbohydrate-binding surface. In addition to the X-ray crystal structures, we explored how the presence of ions affects the stability of the protein, compared the new GbpA LPMO structures to those of other LPMOs, and discussed the relevance of our discovery for bacterial survival. Calcium ions, abundant in natural sources of chitin, have been found to have the strongest effect on GbpA stability. Our findings suggest a *V. cholerae-*specific cation-binding site in GbpA that may fine-tune activity and binding to the different substrates during environmental survival and host infection.

## Introduction

Cholera is an ancient and severe diarrheal disease.^1^ The current cholera pandemic is responsible for over 140.000 deaths annually in communities and countries where proper water sanitation is limited, especially in the wake of natural disasters and armed conflicts.^2^ The Gram-negative bacterium *Vibrio cholerae* is the pathogenic agent responsible for cholera.

*V. cholerae* is a facultative pathogen found in aquatic environments worldwide. Traditionally assumed endemic to tropical regions, it is now regarded as a cosmopolitan species able to survive in a wide range of conditions.^3^ As many other members of the *Vibrio* genus, *V. cholerae* can be found both as free-living bacterium and attached to biotic and abiotic surfaces,^3–5^ where it often forms biofilms.^6,7^ Among the multiple surfaces and organisms that act as environmental reservoirs of *V. cholerae*, chitin and its presence in the exoskeleton of planktonic crustaceans are of particular relevance (Figure 1). Attachment to chitin provides *V. cholerae* with an abundant and stable carbon and nitrogen source, allows lateral transmission of genetic mobile elements linked with pathogenicity^8^ and provides protection from predators.^6,9^ In addition to its role as a reservoir, chitin acts as a critical factor for cholera transmission.^10^ Attachment to chitin is mediated by a range of colonization factors and adhesins.^6^ A colonization factor of particular importance is *N*-acetyl glucosamine (GlcNAc) binding protein A (GbpA),^11^ which is found in all *V. cholerae* strains.^12^

**Figure 1.**
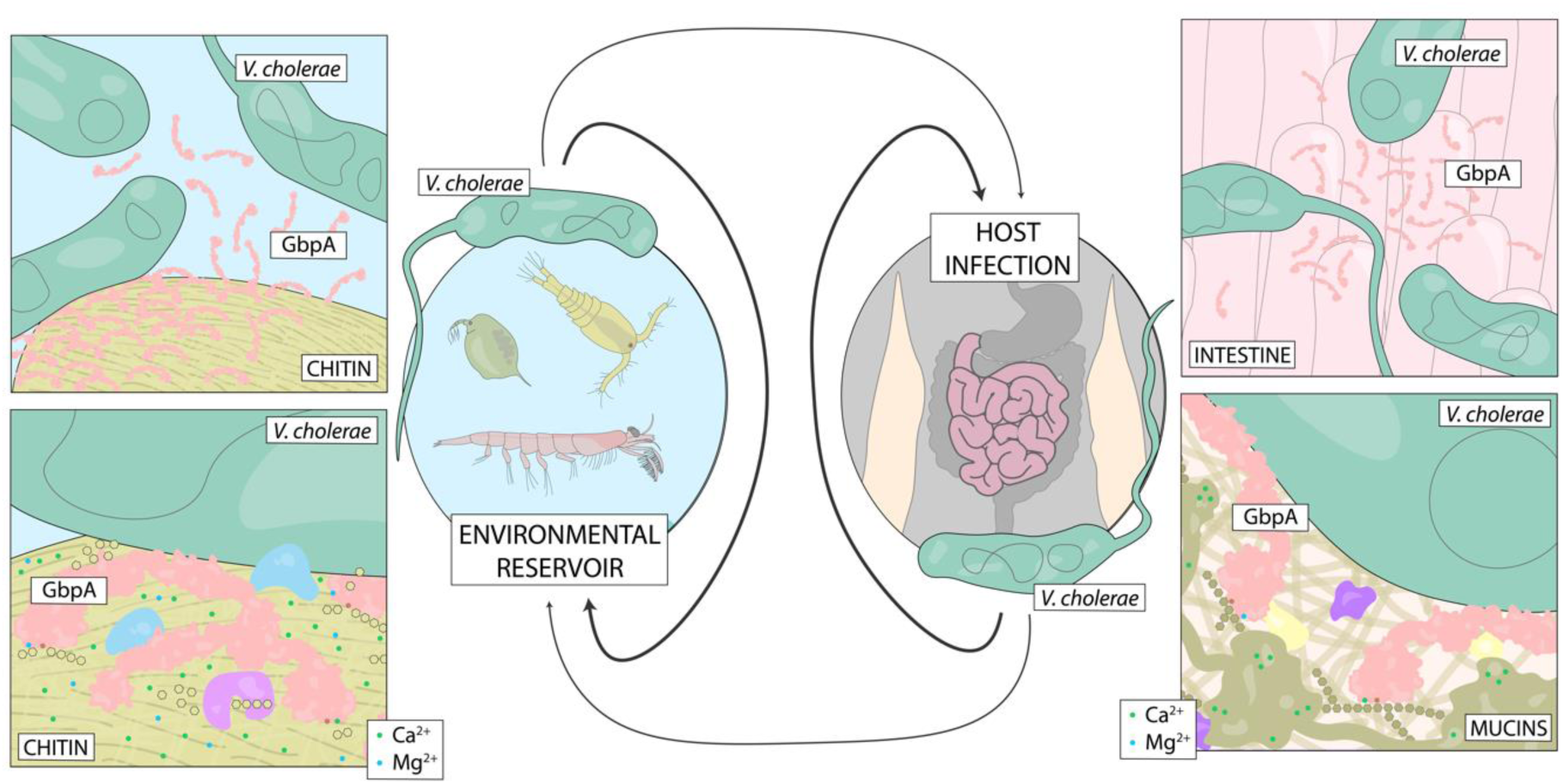
Different niches of *V. cholerae* and functions of GbpA. As a member of the marine microbiome, *V. cholerae* survives attached to biotic and abiotic surfaces. Of particular importance are the chitinous exoskeletons of marine crustaceans found in zooplankton, where *V. cholerae* forms microcolonies and uses the crystalline polysaccharide as a source of nutrients. To this process, GbpA is of particular importance, acting as a colonization factor by binding chitin and as part of the chitin utilization machinery, oxidatively degrading the polysaccharide chains, thereby allowing further processing by other enzymes. During pathogenesis and host infection, *V. cholerae* colonizes the intestine using colonization factors that mediate attachment. Here, GbpA also acts as a colonization factor by recognizing GlcNAc moieties present in the intestine, particularly from the highly glycosylated mucins. Crucially, different concentrations of calcium and magnesium ions are found in the different substrates of GbpA.

GbpA can bind the GlcNAc chains of chitin,^13^ and oxidatively degrades the polymers by means of its lytic polysaccharide monooxygenase (LPMO) activity.^14^ Discovered only recently,^15^ LPMOs are of interest for biomass conversion due to their ability to degrade recalcitrant polysaccharides.^16^ The oxidative activity of LPMOs depends on a copper ion bound by a histidine brace motif. Interestingly, bacterial LPMOs are often associated with pathogenicity^17^ – for instance, CbpD, the tri-modular LPMO of *Pseudomonas aeruginosa*, is crucial for systemic infection.^18,19^ In pathogenic *V. cholerae* strains, GbpA mediates binding to GlcNAc moieties in host intestinal mucins, making it an important virulence factor in addition to its role in environmental survival (Figure 1).^20,21^ The structure of GbpA has been quite well characterized: the crystal structure of the first three domains of the four-domain protein has been solved by X-ray crystallography, showing that domain 3 folds back on LPMO domain 1; whereas the solution structure of the full-length protein determined by small-angle X-ray scattering (SAXS) reveals an elongated conformation.^13^ The first and fourth domains are responsible for chitin binding, with only the first domain showing LPMO activity and additionally binding intestinal mucins.^13^ The second and third domains are likely responsible for interaction with still unidentified bacterial components.^13^

Salinity and the concentrations of specific ions change significantly between the different niches that pathogenic *V. cholerae* colonize, *e.g.*, estuarine waters and the human intestine. Previous studies have investigated the optimal salinity for *V. cholerae* growth, survival and attachment to chitin^22^ as well as the effects of different salts on its attachment to inert surfaces.^4^ Nonetheless, to date, very little is known about how salinity and specific ions influence *V. cholerae* and its ability to colonize and survive on different surfaces, from the environmental reservoirs to the highly complex intestinal tracts of its hosts. Here, we report the discovery of a previously uncharacterized ion binding site close to the active site of the GbpA LPMO domain, which affects protein stability and potentially fine-tunes its oxidative activity and binding to different substrates.

## Results

### New cation binding site identified close to LPMO active site

The LPMO domain of GbpA was saturated with copper, and single crystals suitable for diffraction were obtained in the presence of calcium or potassium ions. The crystal structures of the LPMO-calcium and LPMO-potassium complexes were refined to 1.8 Å and 1.5 Å, respectively, and *R*/*R*_free_ values of 0.179/0.228 and 0.164/0.192 (Table 1). The structures are deposited in the Protein Data Bank (PDB)^23^ with accession IDs 7PB7 (Ca^2+^) and 7PB6 (K^+^). In both structures, a cation binding site was identified close to the copper-binding active site of the LPMO. The nature of the cation was predicted to be the dominant cation species present in the experiments (0.2 M calcium and 0.1 M potassium, respectively), and validated based on geometry analysis, binding distances, electron density peak height and anomalous diffraction analysis (see *Experimental Procedures* for details). The cation binding site is formed mainly by the loop located between strands 7 and 8 of the core β-sandwich characteristic of LPMOs^24^. This includes the backbone carbonyl groups of Val186, Thr189 and Ala191 as well as the side chain carboxylate of Asp185. In addition, coordination by the side chain carboxylate of Asp70, located in the L2 region,^24^ is key to binding (Figure 2B and 2C). The coordination geometry for K^+^ is that of a tetragonal bipyramid, *i.e.*, octahedral (Figure 1C), whereas Ca^2+^ coordination is characterized by a distorted pentagonal bipyramid, with the two carboxyl oxygens of Asp70 involved in the interaction (Figure 2B). The distances as well as the coordination geometries are consistent with the expected values for the corresponding metal ions.^25^ A water molecule completes the coordination sphere of both cations. In particular, a pentagonal bipyramid is typical for divalent calcium, and unusual for many other ions.^26,27^

**Figure 2.**
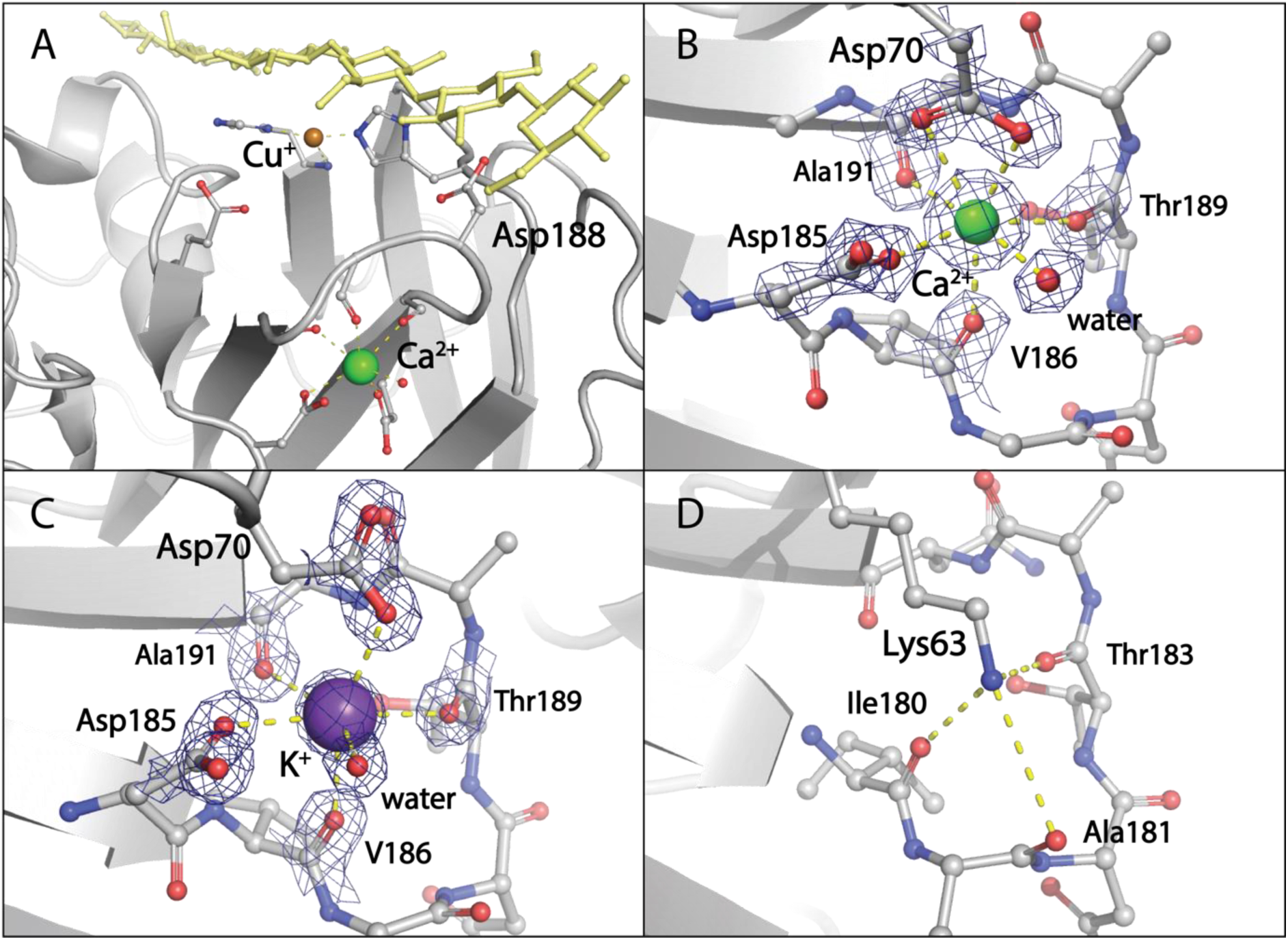
LPMO cation-binding sites. **A**. Overview of the newly identified cation-binding site in proximity of the GbpA active site and carbohydrate-binding surface. Ca^2+^ is represented as green sphere; the active-site copper ion as bronze sphere, bound to histidine-brace motif characteristic of LPMOs. The carbohydrate substrate (yellow) has been manually modeled in its expected position from Tandrup *et al.*^50^, taking into account information from Bissaro *et al.*^28^. Important residues belonging to the active site, the chitin binding surface (in particular Asp188) and the newly identified metal binding site are depicted in stick representation. **B.** Close-up view of cation-binding site featuring Ca^2+^ (green sphere with pentagonal bipyramidal coordination indicated by dashed lines), with sigma-A-weighted 2m*F*o-D*F*c map (blue mesh) contoured at 2σ (PDB ID: 7PB7; this work). **C.** Close-up view of cation-binding site featuring K^+^ (purple sphere with tetragonal bipyramidal coordination), with σ_A_-weighted 2m*F*o-D*F*c map contoured at 2σ (PDB ID: 7PB6; this work). Note the different side chain conformation of Asp70 (at top center of figure panel), which results in different metal ion coordination of K^+^ compared to Ca^2+^ (monovalent instead of divalent interaction). **D.** In other AA10 LPMOs, the position of Asp70 in the L2 loop is occupied by a conserved lysine residue (here: Lys63), which interacts with the backbone of the residues in the loop connecting β-strands 7 and 8 (shown here for *Serratia marcescens* chitin-binding protein 21 (CBP21); PDB ID: 2BEM^51^).

**Table 1.** X-ray data collection and statistics.

**(a) Data collection**
|  | GbpA<sub>LPMO</sub> (Ca<sup>2+</sup>) | GbpA<sub>LPMO</sub> (K<sup>+</sup>) |
| --- | --- | --- |
| Beamline | BM30A | BM14 |
| Wavelength (Å) | 0.9797 | 0.9537 |
| Resolution range | 41.0-1.8 (1.90-1.80) | 46.6-1.5 (1.59-1.50) |
| Space group | *P* 2<sub>1</sub> | *P* 4 |
| Cell parameters |  |  |
| a, b, c (Å) | 40.6, 45.2, 99.6 | 46.6, 46.6, 157.4 |
| α, β, γ (°) | 90, 101.1, 90 | 90, 90, 90 |
| *R*<sub>meas</sub> (%) | 0.197 (0.969) | 0.062 (0.263) |
| CC<sub>1/2</sub> | 0.993 (0.703) | 0.999 (0.973) |
| Mean I/σ | 7.8 (1.8) | 28.4 (7.9) |
| Completeness (%) | 99.5 (98.5) | 98.5 (90.8) |
| Multiplicity | 4.1 (4.1) | 13.4 (10.9) |
| Unique reflections | 33,236 (5267) | 52,428 (7798) |

**(b) Refinement**
|  | GbpA<sub>LPMO</sub> (Ca<sup>2+</sup>) | GbpA<sub>LPMO</sub> (K<sup>+</sup>) |
| --- | --- | --- |
| Resolution (Å) | 34.3 – 1.80 | 40.1 – 1.50 |
| *R*<sub>work</sub> / *R*<sub>free</sub> | 0.179 / 0.228 | 0.164 / 0.192 |
| Macromolecules/a.u. | 2 | 2 |
| Number of non-hydrogen atoms |  |  |
| Protein | 2798 | 2828 |
| Ligands | 4 | 24 |
| Waters | 512 | 565 |
| *B*-factors (Å<sup>2</sup>) |  |  |
| Protein | 14.3 | 12.0 |
| Ligands | 12.4 | 17.9 |
| Waters | 27.0 | 24.5 |
| r.m.s.d. from ideal values |  |  |
| Bond length (Å) | 0.005 | 0.007 |
| Bond angles (deg) | 0.719 | 0.807 |
| Ramachandran Plot |  |  |
| Favored (%) | 96.9 | 97.8 |
| Outliers (%) | 3.1 | 2.2 |
| PDB ID | 7PB7 | 7PB6 |

Since the presence of a cation fixes the loop formed by residues 186-189 to the sequentially distal Asp70 located in the L2 region^24^ (Figure 2B and 2C), we hypothesized that it could confer stabilization of the LPMO domain. Furthermore, the same loop region is part of the LPMO carbohydrate-binding surface, with Asp188 known to be involved in chitin binding (Figure 2A).^28^

### Effect of metal ions on GbpA thermostability

Differential scanning fluorimetry was used to study the effect of different cations on the GbpA LPMO domain (GbpA_LPMO_) and on the full-length protein (GbpA_FL_). A significant stabilizing effect was observed in the presence of calcium, with an increase in melting temperatures (ΔT_m_) of 3.4 °C and 3.8 °C for GbpA_LPMO_ and GbpA_FL_, respectively (Table 2). The analysis was extended to different calcium concentrations (Figure 3), for both apo and copper-bound GbpA_FL_. Here, only apo GbpA showed saturable binding to calcium with a calculated dissociation constant (*K*_d_) of 0.22 mM. To probe if stabilization by calcium is attributed to the identified binding site, Asp70 was substituted with Ala in GbpA_FL_. This GbpA_FL_D70A variant exhibited similar thermal stability as wild-type (WT) GbpA in the absence of calcium, but completely lost the additional stabilizing effect caused by calcium (Figure 3B, Table 2).

**Figure 3.**
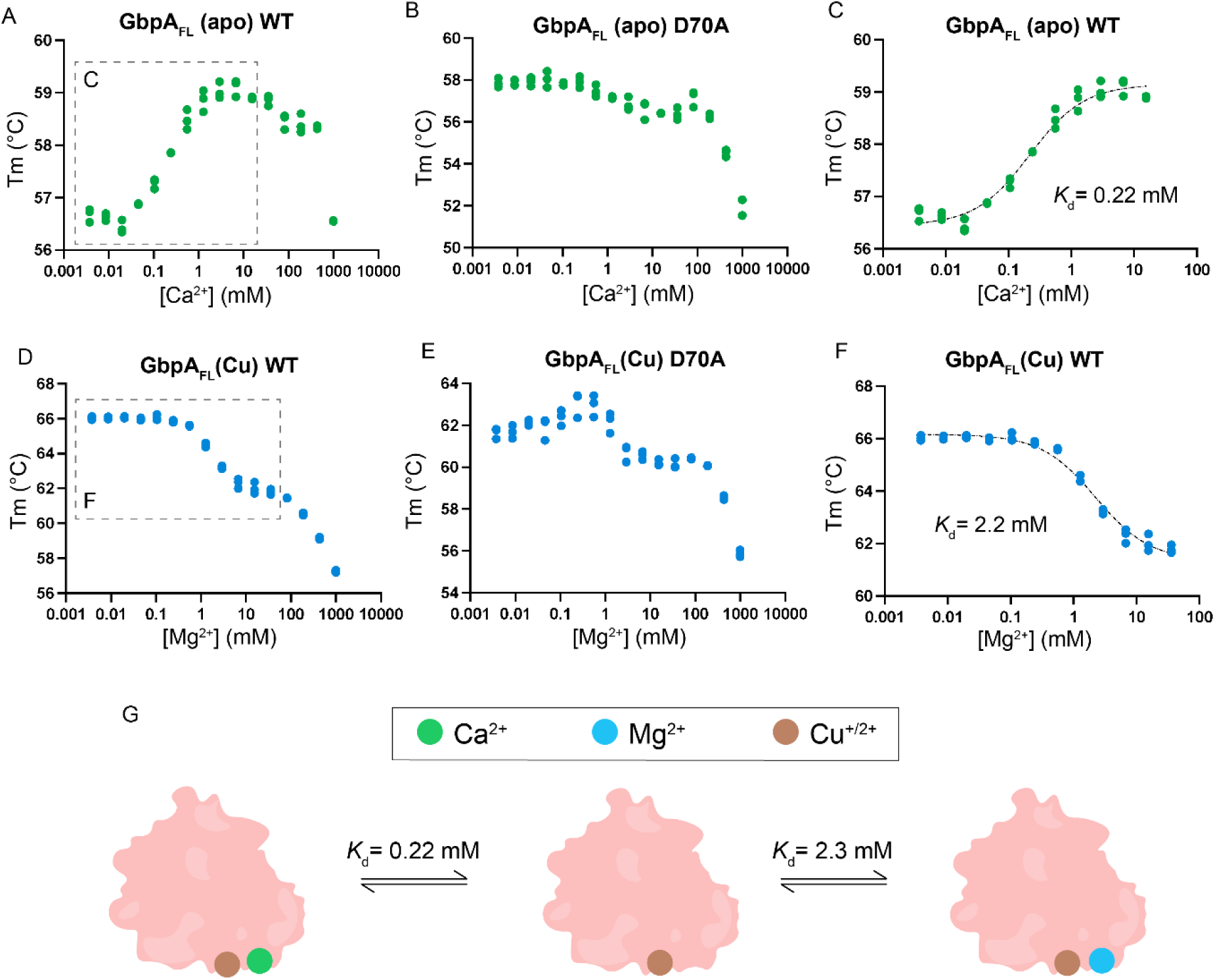
Specific effects of divalent ions on GbpA stability. **A, B** and **C**. Effects of calcium ion concentration on apo GbpA_FL_ stability. Note the specific and saturable stabilization at physiological ion concentrations observed only for wild-type (WT) GbpA. **D, E** and **F.** Effects of magnesium ion concentration on copper-saturated GbpA_FL_. Specific and saturable destabilization is observed for WT GbpA. **G**. Graphical representation of GbpA metal-binding states. Each state potentially has different substrate-binding affinities and catalytic activities. For clarity, only the representation of the GbpA LPMO domain is shown, excluding the second, third and fourth domains. The dissociation constant of copper-bound GbpA for calcium is assumed to be the same as for the apo protein.

**Table 2.** Stability measurements by differential scanning fluorimetry. Effect of different salts on melting temperature of GbpA and GbpA variants (comparison to sample without added salts, top row). For these experiments, the protein was in its apo form, not bound to copper.

|  | GbpA<sub>FL</sub> WT | GbpA<sub>LPMO</sub> WT<sup>b</sup> | GbpA<sub>FL</sub> D70A | GbpA<sub>FL</sub> D70K |
| --- | --- | --- | --- | --- |
| **No salt<sup>a</sup>** | 54.4 ± 0.01 °C | 56.4 °C ± 0.04 °C | 56.8 °C ± 0.09 °C | 54.8 ± 0.24 °C |
| **CaCl<sub>2</sub>** | 3.8 °C | 3.4 °C | -2.0 °C | -0.2 °C |
| **MgCl<sub>2</sub>** | -1.6 °C | -2.1 °C | -0.9 °C | 1.2 °C |
| **KCl** | 1.0 °C | 1.2 °C | 1.2 °C | 1.8 °C |
| **NaCl** | 0.8 °C | 0.9 °C | 1.1 °C | 1.9 °C |

<sup>a</sup> Standard deviations are similar for the experiments with salts. <sup>b</sup>GbpA<sub>LPMO</sub> refers to the isolated LPMO domain, whereas FL refers to full-length GbpA.

When analyzing magnesium binding to GbpA at different ion concentrations, a saturable destabilizing effect was observed for the protein that contained copper in the active site (*K*_d_ = 2.2 mM; Figure 3D and 3F). Magnesium most likely binds to the same site as calcium, since this destabilizing effect is not clearly observed for the GbpA_FL_D70A variant (Figure 3E).

At high concentration, divalent cations appear to cause a general non-specific destabilizing effect, as observed for all magnesium and calcium conditions (Figure 3, Figure S1). In contrast, monovalent cations appear to confer only small stabilizing or destabilizing effects on GbpA, with equivalent trends observed for Na^+^ and K^+^ (Table 2, Figure S1). The changes in stability induced by monovalent and divalent cations do not affect the overall shape of GbpA, as shown by SAXS (Figure S2). The shape of GbpA was similar for all the tested salts (NaCl, KCl, CaCl_2_ and MgCl_2_), with a radius of gyration (Rg) of approximately 38-39 Å.

### The identified cation binding site mimics conserved contacts in related LPMOs

Multiple sequence alignments for GbpA orthologs among the *Vibrio* genus clade show that conservation of the metal-ion binding site is very low, particularly for Asp70. Extending the multiple sequence alignment to more phylogenetically distant LPMOs (classified as AA10 in the CAZy database), we noticed that in 45% of the annotated sequences, the residue found in the position equivalent to Asp70 of GbpA is a lysine. Lysine is also the most common residue in that position for the AA10 LPMO structures deposited in the PDB (Figure 4). Interestingly, the positively charged lysine ammonium group occupies the position of the metal ions and interacts with the backbone residues that constitute the metal-binding site. This is exemplified for chitin binding protein 21 (CBP21) from *Serratia marcescens* in Figure 2D, chosen as a representative example of AA10 LPMO structures featuring a lysine in the L2 region loop. A structural alignment of all the experimentally obtained AA10 LPMO structures with lysine in the position equivalent to Asp70 in GbpA is shown in Figure 5.

**Figure 4.**
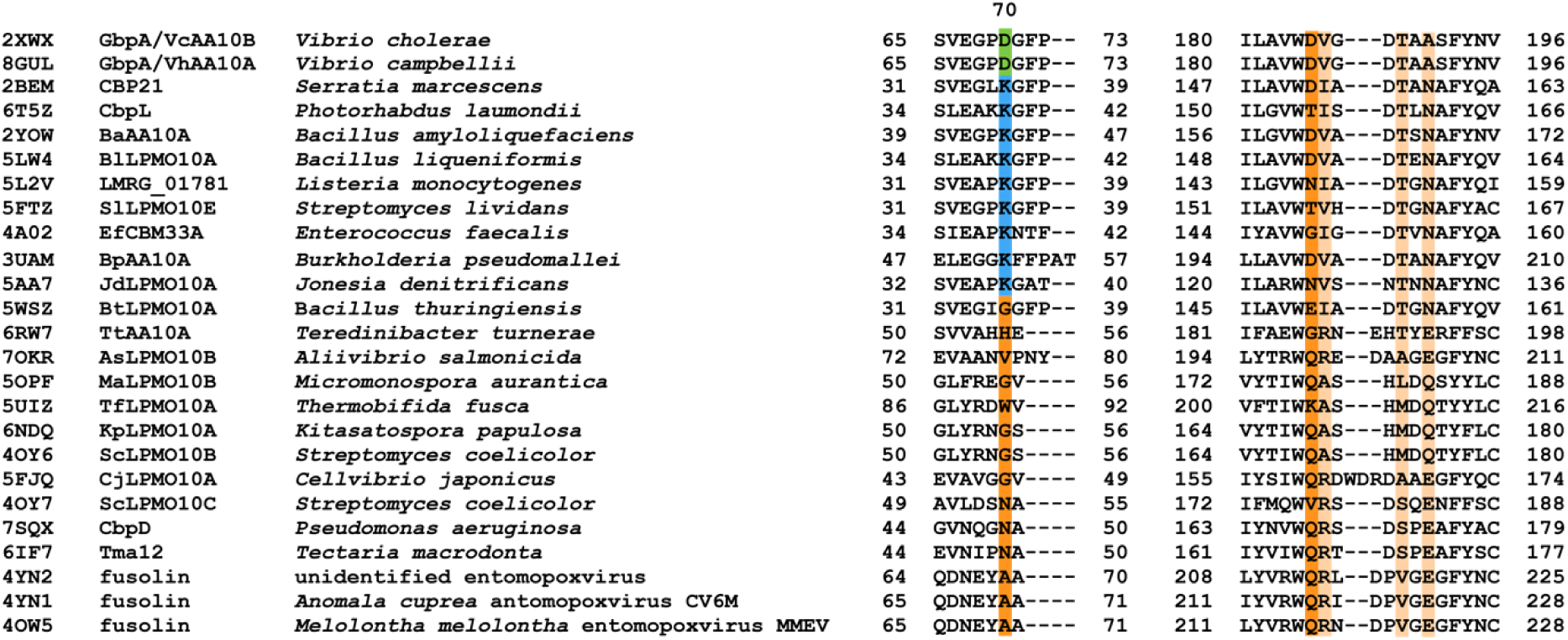
Multiple sequence alignment of AA10 LPMOs. Included in this structure-based alignment are all AA10 LPMO structures currently available in the PDB (with PDB IDs given on the left). Residue numbers are indicated for protein elements involved in cation-binding in GbpA, involving L2 loop residues and β-strands 7 to 8, with residues coordinating the metal ion highlighted. Asp70 (D in one-letter code; highlighted in green) is often replaced by Lys (K in one-letter code; highlighted in blue) in other AA10 LPMOs.

**Figure 5.**
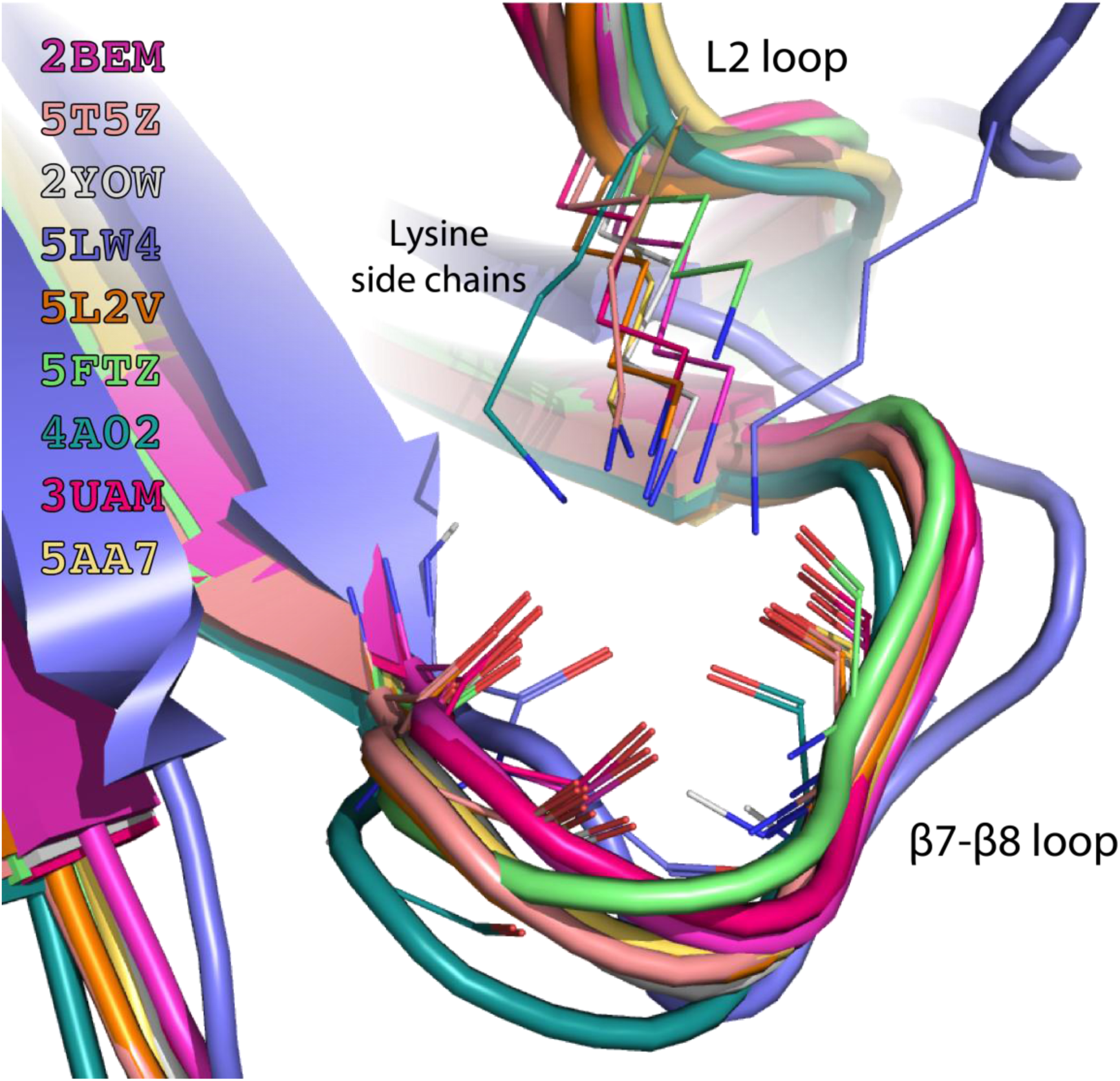
Structural alignment of AA10 LPMOs. The alignment includes all AA10 LPMO structures currently available in the PDB that exhibit a lysine residue in the L2 loop. The structures are color-coded, and interactions with the β7-β8 loop are shown. PDB IDs are given to the left. *Serratia marcescens* chitin-binding protein 21 (CBP21; PDB ID: 2BEM^51^) is represented in magenta, with detailed structural interactions of L2 loop residue Lys63 that stabilize the loop between β-strands 7 and 8 depicted in Figure 2D.

In order to better understand the effect that the emergence of the binding site might have had for GbpA, we engineered an additional GbpA_FL_ variant, D70K. As expected, this substitution eliminated the specific stabilizing effect by calcium (Table 2). However, we also did not observe any significant compensating effect in form of general protein stabilization in the absence of salts (ΔT_m_ = 0.4 °C, shown in Table 2). Therefore, the presence of the positively charged ammonium group in the metal binding site did not mimic the binding of calcium, but is more similar to monovalent ions like potassium, which has little effect on GbpA stability (Table 2, Figure S1).

## Discussion

Here, we report the discovery of a metal-binding site in GbpA close to the LPMO active site. We determined crystal structures of the GbpA LPMO domain in complex with either calcium or potassium ions to high resolution, and tested the effect of different metal ions on GbpA thermostability. Calcium and magnesium were found to have specific effects on GbpA stability, with calcium stabilizing and magnesium destabilizing the protein. Stabilization by calcium (in the high-micromolar and low-millimolar range) was only observed for GbpA in the apo state, and not for the copper-saturated protein. However, the crystal structure determined by us (PDB ID: 7PB7) clearly shows that calcium can bind to GbpA also in the presence of copper. We therefore assume that the stabilizing effect of calcium is likely shielded by the strongly stabilizing copper ion, which is reported to bind to the histidine brace with high affinity, in the low nanomolar range.^29^ In contrast, magnesium was found to destabilize GbpA (in the low-millimolar range), which could imply an increase in flexibility that may be required for chitin processing. It is therefore likely that calcium and magnesium ions have a biological role, regulating GbpA binding and activity.

How could these ions achieve this? GbpA is known to bind chitin through domains 1 and 4^13^, whereas binding to mucins appears to only involve the LPMO domain 1,^13^ however, relatively little detail is known about these interactions at the molecular level.^30^ An additional challenge is assessing what ionic compositions *V. cholerae* might be exposed to in its local microenvironment during environmental survival and intestinal colonization is a complex task.

Calcium and magnesium concentrations in aquatic environments are generally low, and limited by the solubility of their carbonate salts^31^ to around 0.13 mM for calcium and 1.4 mM for magnesium ions (both measured at 25 °C). Interestingly, the dissociation constants obtained for both salts are approximately twice their solubility in the ocean, with *K*_d_ = 0.22 mM and 2.2 mM for calcium and magnesium ions, respectively. Although dissociation constants calculated by thermal shift analysis are obtained at the non-physiological melting temperature of the protein and could thus differ significantly from the effective values, it seems that GbpA would require to be exposed to higher ion concentrations than those generally present in the marine environment. However, the chitinous exoskeleton of crustaceans is mineralized mostly with calcium and – in smaller amounts – magnesium amorphous precipitates.^32^ Therefore, GbpA is most likely exposed to significantly higher local ion concentrations when binding to chitin of crustaceans. Furthermore, the amount of mineralization and the nanostructure of the ionic precipitate varies widely for different body parts^33^ as well as among different species of crustaceans. It is thus plausible that GbpA could establish preferential binding to certain surfaces or organisms, paving the way towards colony and biofilm formation. And the interplay between calcium and magnesium may facilitate binding and release during chitin processing.

Conversely, in the mammalian gut, where GbpA is involved in pathogenesis, total calcium and magnesium levels are likely to be lower than in chitin. Concentrations of these cations in the human serum are 2.2–2.6 mM for calcium and 0.7–1.1 mM for magnesium,^34^ however, this does not translate directly to the concentrations in the gut. Ion concentrations are difficult to estimate due to the highly complex environment and composition of the organ, where local ion concentrations may be variable and dependent on factors such as diet and absorption by the host,^35^ as well as interactions with the native microbiota. Nevertheless, calcium concentrations in the gut are sufficiently high to be relevant for the conformation and mechanical properties of mucins.^36,37^ Mucins contain multiple calcium-binding sites in their von Willebrand factor (vWF) and CysD domains and expand in the absence of these ions.^38^ We hypothesize that calcium binding to GbpA could represent a competition mechanism by which the presence of GbpA in the gut could affect mucin conformations, aiding colonization by *V. cholerae* through a less dense mucin structure and possibly facilitating toxin entry.

The role of K^+^ remains enigmatic. In terms of protein stability, the presence of a potassium ion in the metal binding site might be equivalent to that of the lysine residue from loop L2, which occupies the metal binding site in some AA10 LPMOs. However, the relatively conserved nature of the lysine could indicate an important role in other AA10 LPMOs. We hypothesize that monovalent ions, generally more soluble and abundant,^39^ could represent opportunistic binders in the absence of calcium or magnesium and thus participate in a regulation mechanism by ion exchange when calcium or magnesium concentrations are below a certain threshold.

Information obtained from activity and chitin binding measurements for GbpA in the presence of different salts would be highly relevant. However, such experiments are challenging given that assays on LPMO activity of GbpA are done on chitin fibers from natural origins, where controlling the presence and amounts of specific ions may prove difficult. It may also be interesting to extend the bioinformatics studies to search for clues to the relevance of the binding site and investigate if it is present in LPMOs from phylogenetically more distant organisms.

While the metal binding site is rare among orthologs of GbpA from the *Vibrio* genus, it is conserved among all *V. cholerae* strains analyzed by Strauder *et al.*^12^, both from environmental and pathogenic origins. This suggests that the binding site is likely an important adaptation to the lifestyle of these bacteria, whereas other species, featuring a conserved lysine at this position, lack such a regulation mechanism. This supports the hypothesis that the local structure and organization of the region involving the L2 loop as well as the loop connecting strands 7 and 8 is important for these LPMOs, with potential implications for binding and catalysis.

Summarizing, we report the discovery of a second cation-binding site in GbpA, which is in close proximity to active site and substrate-binding site. We show that calcium and magnesium specifically affect GbpA stability, suggesting that ion binding may modulate binding and catalytic activity also *in vivo*, with potentially important consequences for *V. cholerae*’s survival in different niches, including environmental survival and infectivity.

## Experimental procedures

### Protein expression constructs and mutagenesis

The GbpA gene (UniProt ID: Q9KLD5, residues 24-485) was codon-optimized and cloned into a pET-26b vector by GenScript® between the restriction sites *Nco*I and *Xho*I. The gene was cloned from residue 24 —omitting the natural export signal— so that the PelB leader sequence in the vector would direct the expressed protein for post-translational translocation to the periplasmic space. After translocation, the PelB sequence is cleaved by the expression system. The C-terminal His-tag present in the vector was omitted by the inclusion of a stop codon at the end of the insert. The pET-26b vector contains a kanamycin resistance gene, which we exploited for selection in growth and expression phases. For the expression of the LPMO domain of GbpA the construct is equivalent to the full length GbpA containing the residues 24 to 203.

Point mutations were introduced in the GbpA sequence using the NEB Q5 Site-Directed Mutagenesis Kit. Successful production of mutants was confirmed by sequencing the newly generated vectors.

### Protein production and purification

We used the same expression and purification protocols for producing full-length GbpA, the GbpA LPMO domain and GbpA variants ^40^. Briefly, a 2 mL preculture in generic LB media was grown at 37 °C for 6 hours in presence of 50 μg/mL of kanamycin sulphate in a 17x100 mm culture tube at 220 rpm. 200 μL of the preculture were used to inoculate 25 mL of M9glyc+ media with 50 μg/mL of kanamycin sulphate in a baffled Erlenmeyer flask. 1 L of M9glyc+ media contains 19 g K_2_HPO_4_, 5 g KH_2_PO_4_, 9 g Na_2_HPO_4_, 2.4 g K_2_SO_4_, 5.0 g NH_4_Cl, 18 g glycerol, 10 mM MgCl_2_, 1x MEM vitamins and 1x Trace element solution. 0.1 L of Trace element solution 100x contain 0.6 g FeSO_4_·7H_2_O, 0.6 g CaCl_2_·2H_2_O, 0.12 g MnCl_2_·4H_2_O, 0.08 g CoCl_2_·6H_2_O, 0.07 g ZnSO_4_·7H_2_O, 0.03 g CuCl_2_·2H_2_O, 0.02 g H_3_BO_4_, 0.025 g (NH4)_6_Mo_7_O_24_·4H_2_O and 17 mM EDTA. The culture was grown at 37 °C and 120 rpm. After 16 hours 225 mL of fresh M9glyc+ media with 50 μg/mL of kanamycin sulphate were added to the baffled flask. Around 2 hours later, when the OD_600_ was between 2 and 3, protein production was induced by adding isopropyl β-D-1-thiogalactopyranoside (IPTG) to a concentration of 1 mM, and lowering the temperature to 20 °C. Harvesting was performed after 20 hours by centrifugation at 10,000 rcf for 30 min at 4 °C.

An osmotic shock protocol was used in order to separate the periplasmatic fraction of the collected cells. First, the cell pellet was resuspended in around 5 mL per gram of cell paste of a buffered hypertonic solution (Tris-HCl 20 mM pH 8.0, 25% w/v of D-saccharose and 5 mM ethylenediaminetetraacetic acid (EDTA)). The solution was incubated under gentle stirring for 30 min at 4 °C and centrifuged at 10,000 rcf for 30 min at 4 °C. The supernatant was kept for further purification and the pellet was resuspended in a hypotonic solution (Tris-HCl 20 mM pH 8.0, 5 mM MgCl_2_, 1 mM phenylmethylsulfonyl fluoride (PMSF) and 0.25 mg/mL of lysozyme from chicken egg white (Sigma-Aldrich)). The solution was incubated again for 30 min under gentle stirring at 4 °C and centrifuged at 10,000 rcf for 30 min at 4 °C. The supernatant was also saved for further purification.

The two supernatant fractions were loaded into a HiTrap Q HP 5mL anion exchange column equilibrated in 20 mM Tris-HCl pH 8.0, 50 mM NaCl and eluted with a gradient of 20 column volumes up to 400 mM NaCl. The peaks containing the desired protein were concentrated and loaded into a Superdex 75 10/300 Increase equilibrated in 20 mM Tris-HCl pH 8.0 for a final purification step.

### Multiple sequence alignments

Multiple sequence alignment sequences for the related orthologs in the *Vibrio* species clade were obtained from OrthoDB^41^ (group 52685at662 at *Vibrio* level), while the sequence files for all the annotated members of the AA10 family were obtained from Genbank^42^ based on the annotations in the CAZy^43^ database. Multiple sequence alignments were performed with Clustal Omega^44^ with default parameters and manually inspected in Jalview.^45^

### Differential scanning fluorimetry

Differential scanning fluorimetry (DSF) experiments were carried out in a Prometheus NT.48 system from NanoTemper (NanoDSF), measuring the change in fluorescence by the protein aromatic residues upon thermal denaturation. The samples were buffer exchanged to 20 mM HEPES pH 7.0 with the corresponding salt present at 50 mM with overnight dialysis using Slide-A-Lyzer dialysis cassettes from ThermoFisher scientific. The pH adjustment of the HEPES buffer was done with NaOH to a final concentration of 7mM Na^+^ for the working HEPES concentration of 20 mM. Measurements were done in triplicates for GbpA_FL_WT, GbpA_FL_D70A and GbpA_FL_D70K, whereas GbpA_LPMO_WT was measured in duplicates. The obtained values for each sample type were averaged, with standard deviation between equivalent measurements being lower than 0.3 °C in all cases.

NanoDSF analysis at varying salt concentrations were carried out under the same buffer conditions, 20 mM HEPES pH 7.0. Copper saturation was performed with a 3:1 molar excess of CuCl_2_ followed by incubation for 30 min at room temperature. Unbound copper was then eliminated through buffer exchange. Measurements were done in triplicates in the conditions for which affinity constants could be calculated. Data analysis, fitting and plotting was performed on using GraphPad Prism. The binding curves were fitted following a single site ligand binding model as described by Vivoli *et al.*^46^

### Small-angle X-ray scattering

SAXS data were collected at the BM29 BioSAXS beamline^47^ at the European Synchrotron Radiation Facility (ESRF) in Grenoble, France. A wavelength λ of 0.992 Å was used, and scattering intensities I(q) were recorded as a function of the scattering vector q = (4π/λ) sin(θ), where 2θ is the scattering angle. The acquisition was done in the q-range 0.0037 to 0.5 Å^-1^. Ten frames were collected per sample and 20 for each buffer. Frames with radiation damage were discarded, the rest were used for further processing.

The DOI associated with the data collection is 10.15151/ESRF-ES-649173298. Samples were measured in 50 mM Bis-Tris propane pH 7 and 500 mM of the respective salts at 37 °C, data with 10 mM NaCl were also recorded for comparison. However, as GbpA was particularly susceptible to radiation damage in the presence of CaCl_2_, parameters were optimized to limit this issue, necessitating a HEPES buffer at 20 mM and pH 7.0, with the CaCl_2_ present at only 50 mM, and attenuation of the beam. Statistics for the CaCl_2_ data are hence weaker. The scattering contribution of the buffer was subtracted from measurements and the data were calibrated to absolute scale using the scattering of H_2_O as a standard. Data were collected for GbpA at different concentrations (0.125, 0.25, 0.5, 1, 2 and 4 mg/mL), those datasets were then merged with *ALMERGE*^48^ from the *ATSAS* package^49^, generating datasets extrapolated to zero solute concentration. R_g_ values were obtained through Guinier analysis using home-written MATLAB scripts. I(q) vs. q plots and dimensionless Kratky plots were made with MATLAB.

### Protein crystallization and X-ray data collection and refinement

Before crystallization of the LPMO domain, the protein was treated with a 3:1 molar excess of CuCl_2_ and incubated for 30 min at room temperature, then desalted and concentrated to 20 mg/mL. Crystals of good diffraction quality were obtained from 0.2 M calcium chloride dihydrate, 0.1 M HEPES pH 7.0, 20% w/v PEG6000 or 0.1 M potassium thiocyanate, 30% PEG 2000 monomethyl ether (MME), respectively. Diffraction data extending to 1.8 Å were collected at beam line BM30 at the ESRF (Grenoble, France) from the crystals obtained from the calcium-containing condition. X-ray data were auto-processed at the ESRF by the *EDNA* pipeline^82^. The structure was phased by molecular replacement (MR) with the *PHENIX* crystallographic software package^83^, using domain I of the GbpA structure (PDB ID:2XWX; DI-DIII)^13^ as a search model, and refined in alternating cycles of manual model building and refinement with *PHENIX*^83^ and *Coot*^84^. Water molecules were added at late stages of the refinement, initially using the automated water picking routine of *PHENIX*.^83^ These sites were then inspected individually. Strong globular electron density was found in the proposed binding site and calcium built into it. While a water molecule could not explain the strong electron density, leaving a residual difference density peak in the binding site, the calcium ion matched the electron density very well. Moreover, coordination geometry and bond distances to neighboring residues and water molecules further strengthened the interpretation as a calcium site.

For crystals obtained from the potassium-containing condition, diffraction data extending to 1.5 Å were collected at ESRF beamline BM14. To aid structural analysis, in particular of the cation-binding site, we collected a complete dataset supporting anomalous signal, and further cut the data to only contain strong anomalous data. The structure was phased with MR, modeled and refined similarly to the calcium-bound structure. Strong density was found in the same site as for calcium. The identity of potassium over other ions or molecules, such as sodium or water, was confirmed by the presence of strong anomalous signal. Furthermore, bond distances to local environment supported this identity. Statistics on data collection and refinement can be found in Table 1, and the final models were deposited in the PDB with accession codes 7PB6 (potassium-GbpA_LPMO_) and 7PB7 (calcium-GbpA_LPMO_).

## Acknowledgements

We would like to thank Joan Montserrat Canals for his help retrieving the sequences for AA10 LPMOs from the CAZy database, and Salvador Almagro-Moreno for interesting discussions and comments on the manuscript. X-ray scattering experiments were performed on beamlines BM29, BM14 and BM30A at the European Synchrotron Radiation Facility (ESRF), Grenoble, France. We are grateful to Local Contacts Dihia Moussaoui, Gabriele Giachin, Martha Brennich, Hassan Belrhali and Franck Borel at the ESRF for providing assistance in using the beamlines. Differential scanning fluorimetry was carried out Helse Sør-Øst Regional Core Facility for Structural Biology. All other work was performed at the UiO Structural Biology core facilities and RECX.

## Author contributions

K.B.-A. conceived the study. K.B.-A. and M.M.-C. designed the genetic constructs (M.M.-C. those of the GbpA variants); M.M.-C., K.B-A. and H.V.S. produced and purified the proteins. G.V.-K. supported the project with expertise and suggestions. M.M.-C. performed the multiple-sequence alignments, guided by G.C.. M.M.-C. and H.V.S. carried out the nanoDSF experiments (analyzed by M.M.-C.), and H.V.S. the SAXS experiments. K.B.-A. crystallized the protein, and determined and refined the crystal structures, which were additionally validated by U.K.. M.M.-C., K,B.-A. and U.K. analyzed the structural data. K.B.-A. supervised H.V.S., and U.K. supervised M.M.-C., K.B.-A. and H.V.S.. M.M.-C. wrote the first draft of the manuscript, which was revised together with K. B.-A. and U.K. based on input from all authors.

---

**Figure S1.**
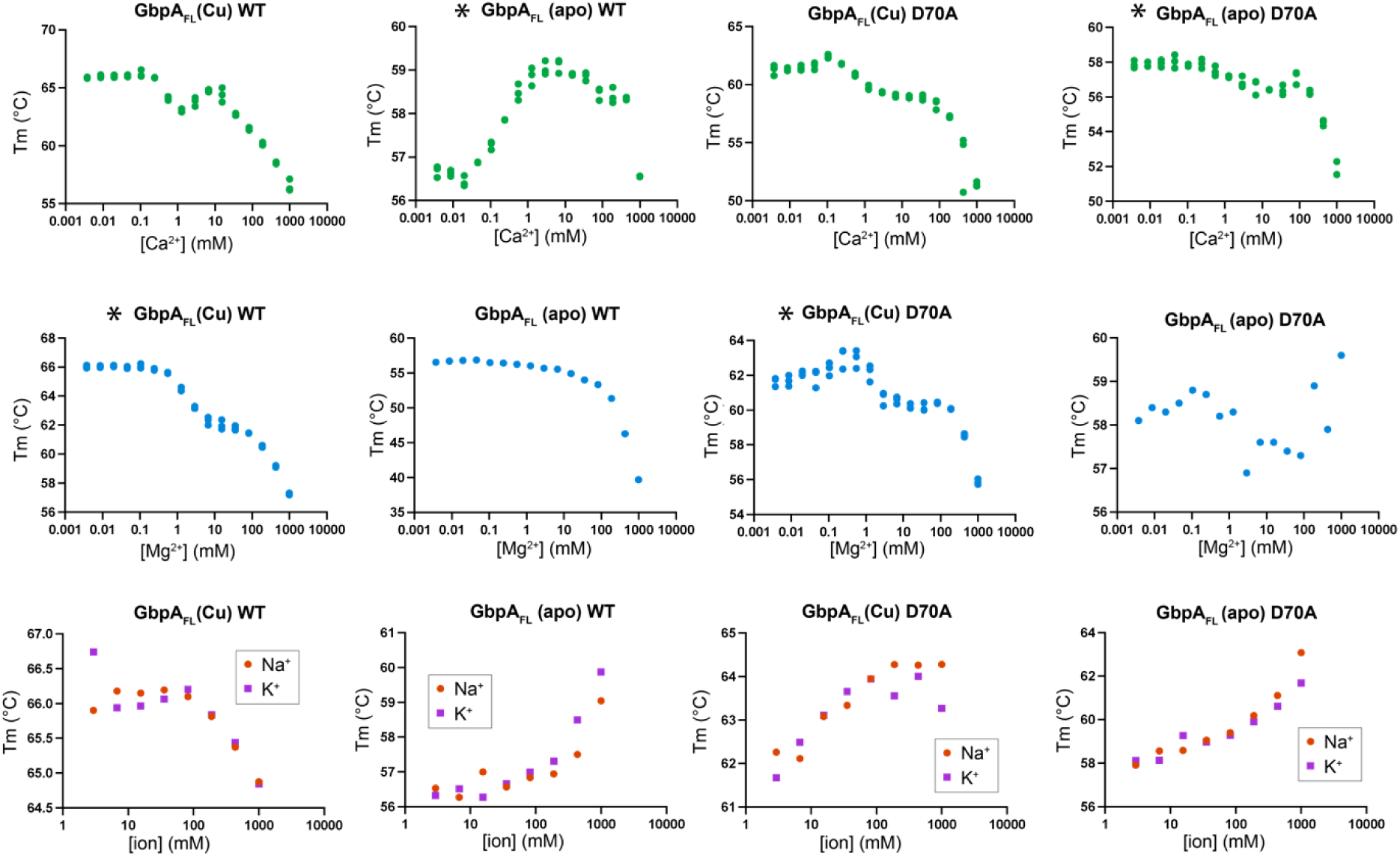
Effects of different cations on GbpA stability. Stability of GbpA_FL_ subjected to different cations and ion concentrations. Experiments were performed for GbpA WT and D70A variants, both for apo (apo) and copper-saturated protein (Cu). Note the general destabilizing trend for divalent cations and stabilization by monovalent ions. Panels with a star (*) are also part of Figure 2.

**Figure S2.**
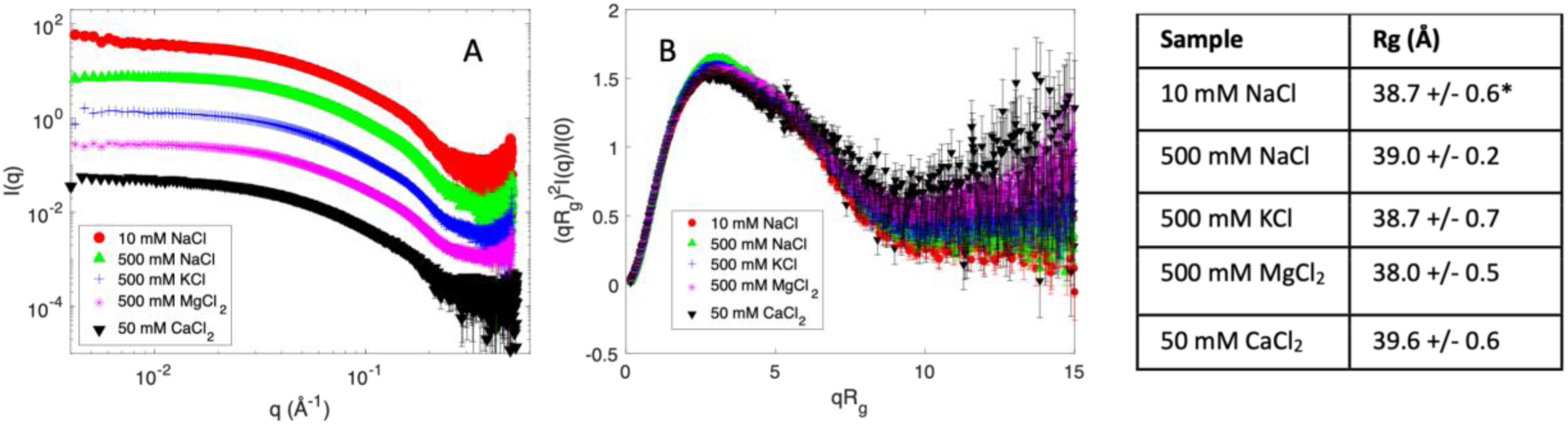
SAXS data for GbpA in the presence of different cations. **A**. Intensities are plotted against the scattering vector q and arbitrarily scaled for visibility. The plots are merged from datasets of concentration series of GbpA exposed to the respective salts, by extrapolating to infinite dilution, hence eliminating concentration-dependent structure factor effects. The plots show limited salt effects on the overall structure of GbpA, but with slight aggregation in 10 mM NaCl, where the salt concentration might be insufficient to screen out inter-particle effects. All data were collected at pH 7.0. For CaCl_2_, data were only collected at 50 mM, and a different buffer (HEPES) was used, since higher salt concentrations and different buffers led to radiation damage. **B**. Dimensionless Kratky plots show that GbpA is folded and globular under all conditions, as the plots are bell-shaped rather than having a consistent inclination at high qR_g_, a characteristic of unfolded proteins. The 500 mM MgCl_2_ and 50 mM CaCl_2_ samples do not reach the same low plateau, but the difference is within the margin of error. In the table to the right, the R_g_ from Guinier approximation is shown for each sample. No significant difference in R_g_ is observed. *For 10 mM NaCl, data below q = 0.012 Å^-1^ were omitted from the Guinier analysis.

**Table S1.**
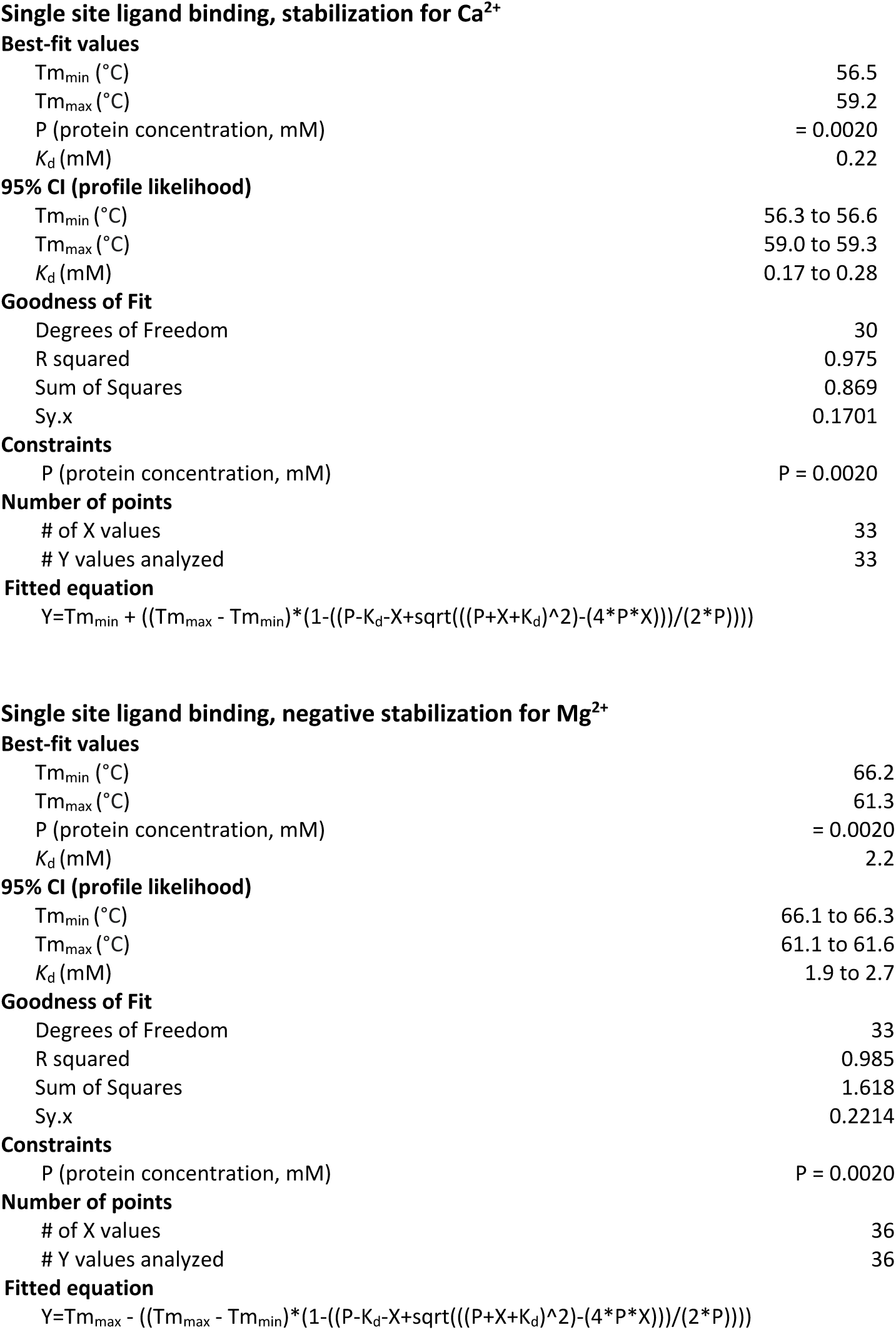
Fitting of dissociation constants (*K*_d_).

**Best-fit values**
|  |  |
| --- | --- |
| $T_{m_{\min}}$ ($^{\circ}\text{C}$) | 56.5 |
| $T_{m_{\max}}$ ($^{\circ}\text{C}$) | 59.2 |
| P (protein concentration, mM) | = 0.0020 |
| $K_d$ (mM) | 0.22 |
| **95% CI (profile likelihood)** |  |
| $T_{m_{\min}}$ ($^{\circ}\text{C}$) | 56.3 to 56.6 |
| $T_{m_{\max}}$ ($^{\circ}\text{C}$) | 59.0 to 59.3 |
| $K_d$ (mM) | 0.17 to 0.28 |
| **Goodness of Fit** |  |
| Degrees of Freedom | 30 |
| R squared | 0.975 |
| Sum of Squares | 0.869 |
| Sy.x | 0.1701 |
| **Constraints** |  |
| P (protein concentration, mM) | P = 0.0020 |
| **Number of points** |  |
| # of X values | 33 |
| # Y values analyzed | 33 |
| **Best-fit values** |  |
| $T_{m_{\min}}$ ($^{\circ}\text{C}$) | 66.2 |
| $T_{m_{\max}}$ ($^{\circ}\text{C}$) | 61.3 |
| P (protein concentration, mM) | = 0.0020 |
| $K_d$ (mM) | 2.2 |
| **95% CI (profile likelihood)** |  |
| $T_{m_{\min}}$ ($^{\circ}\text{C}$) | 66.1 to 66.3 |
| $T_{m_{\max}}$ ($^{\circ}\text{C}$) | 61.1 to 61.6 |
| $K_d$ (mM) | 1.9 to 2.7 |
| **Goodness of Fit** |  |
| Degrees of Freedom | 33 |
| R squared | 0.985 |
| Sum of Squares | 1.618 |
| Sy.x | 0.2214 |
| **Constraints** |  |
| P (protein concentration, mM) | P = 0.0020 |
| **Number of points** |  |
| # of X values | 36 |
| # Y values analyzed | 36 |

